# Stem cell-derived mouse embryos develop within an extra-embryonic yolk sac to form anterior brain regions and a beating heart

**DOI:** 10.1101/2022.08.01.502375

**Authors:** Gianluca Amadei, Charlotte E Handford, Joachim De Jonghe, Florian Hollfelder, David Glover, Magdalena Zernicka-Goetz

**Affiliations:** Department of Physiology, Development and Neuroscience, University of Cambridge, Downing Street, Cambridge CB2 3EG, UK; California Institute of Technology, Division of Biology and Biological Engineering, 1200 E. California Boulevard, Pasadena, CA 91125, USA; Centre for Trophoblast Research, University of Cambridge, Downing Site, Cambridge CB2 3EG, UK; Department of Biochemistry, University of Cambridge, Cambridge, CB2 1GA, UK

## Abstract

Embryo-like structures generated from stem cells can achieve varying developmental milestones, but none have been shown to progress through gastrulation, neurulation, and organogenesis.^1–7^ Here, we show that “ETiX” mouse embryos, established from embryonic stem cells aggregated with trophoblast stem cells and inducible extraembryonic endoderm stem cells, can develop through gastrulation and beyond to undertake neural induction and generate the progenitors needed to create the entire organism. The head-folds of ETiX embryos show anterior expression of Otx2, defining forebrain and midbrain regions that resemble those of the natural mouse embryo. ETiX embryos also develop beating hearts, trunk structures comprising a neural tube and somites, tail buds containing neuromesodermal progenitors and primordial germ cells, and gut tubes derived from definitive endoderm. A fraction of ETiX embryos show neural tube abnormalities, which can be partially rescued by treatment with the metabolically active form of folic acid, reminiscent of common birth defects and therapies in humans. Notably, ETiX embryos also develop a yolk sac with blood islands. Overall, ETiX embryos uniquely recapitulate natural embryos, developing further than any other stem-cell derived model, through multiple post-implantation stages and within extra-embryonic membranes.

---

In natural development, the zygote develops into three initial lineages: the epiblast, which will form the organism, the extraembryonic visceral endoderm (VE), which contributes to the yolk sac, and extraembryonic ectoderm (ExE), which contributes to the placenta. Stem cells derived from these lineages offer the possibility to completely regenerate the mammalian organism from multiple components instead of from a single totipotent zygote.

Embryonic stem cells (ESCs), which are derived from the epiblast, show a remarkable ability to form embryo like-structures upon aggregation. For instance, ESC aggregates (“gastruloids”) embedded in Matrigel can form trunk-like structures with somites, a neural tube and a gut ^1–5^. Although anterior neural development can be promoted in gastruloids by inhibiting the initial burst of WNT activity, they do not faithfully represent the anatomy of natural embryos^6^. Another model, “embryoids” generated from naïve ESCs aggregated with an ectopic morphogen signalling centre, can develop a posterior mid-brain, neural tube, cardiac tissue and the gut tube ^7^. These models form some epiblast structures *de novo*, but do not unlike natural epiblasts they do not undergo gastrulation.

Signals originating from extraembryonic tissues are essential to pattern the pluripotent epiblast and drive the establishment of the anterior-posterior axis. Guided by this requirement for extraembryonic cells in natural development, we have generated synthetic embryos by aggregating ESCs with Trophoblast Stem Cells (TSCs) derived from the precursor of ExE and eXtra-Embryonic eNdoderm (XEN) stem cells derived from the precursors of VE^8^. Substituting XEN cells with “inducible XEN” (iXEN) cells, namely ESCs that transiently express the VE master regulator Gata4, improved the efficiency and developmental potential of these synthetic embryos^9^. Importantly, like natural embryos, these “ETiX” embryos (formerly termed iETX)^9^ specify a migrating anterior visceral endoderm (AVE) and undergo gastrulation movements, which are essential for subsequent stages of development.

A limitation of all of the above strategies has been the failure to accurately recapitulate the development of a natural embryo, that undergoes epithelial-to-mesenchymal transition, gastrulation, organogenesis and the development of anterior brain structures. Here, we identify conditions that allow ETiX embryos to recapitulate gastrulation and natural post-implantation development to embryonic day 8-8.5. Such ETiX embryos possess forebrain and midbrain regions, an organised neural tube, a beating heart and a gut tube. The neural tube is flanked by developing somites and we observe formation of primordial germ cells in the tail region. These structures of the epiblast lie within an extra-embryonic yolk sac that initiates blood island development. To our knowledge, ETiX embryos show the greatest developmental potential and physiological relevance of any stem-cell derived system to date.

## Development of ETiX embryos through neurulation

To identify conditions that support the development of anterior brain structures in ETiX embryos, we seeded ESCs, TSCs and iXEN cells as demonstrated previously^9^ (Fig. 1a), selected ETiX structures that had formed the correct post-implantation morphology on day 4 and moved them to suspension culture. We then took ETiX day 5 structures that had developed a proamniotic cavity, a fully migrated AVE, and were exhibiting gastrulation onset on the opposite side to culture them under conditions supporting the development of natural embryos *ex utero* beyond E5.5^10^. On day 7, we supplemented the medium with glucose and transferred ETiX embryos to rotating culture bottles for one additional day.

**Figure 1:**
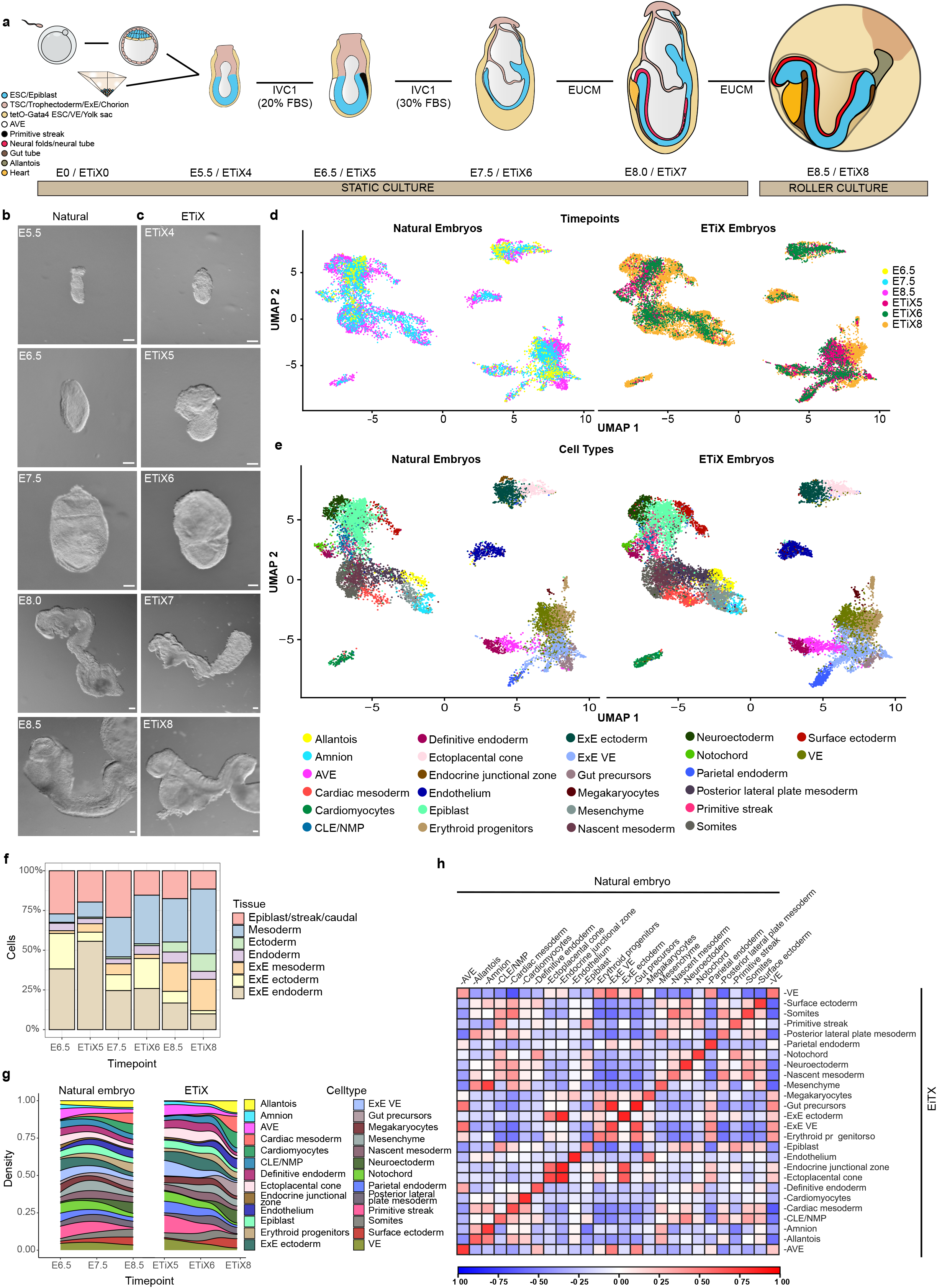
ETiX embryos recapitulate developmental milestones of the natural mouse embryo up to E8.5. **a.** Schematic of ETiX-embryo formation. ETiX embryos are formed by aggregating ESC, TSC and ESC transiently expressing Gata4 and by day 4 generate structures that resemble implantation stage natural E5.5 embryos. They subsequently develop to gastrulation (E6.5/ETiX5) and neurulation (E8.0/ETiX7) before initiating organogenesis (E8.5/ETiX8). Legend: TE: Trophectoderm, Epi: Epiblast, PrE: Primitive Endoderm, ExE: Extraembryonic Ectoderm, VE: Visceral Endoderm, PS: Primitive Streak, AVE: Anterior Visceral Endoderm, HF: Head Fold, H: Heart, G: Gut, T: Tail, Am: Amnion, Al: Allantois, Ch: Chorion, YS: Yolk Sac **b.** Brightfield images of natural mouse embryos and (**c.**) ETiX at different timepoints highlighting their morphological similarities. Scale bar for b,c = 100 μm. **d.** UMAP analysis of single cell RNA-seq data at indicated timepoints for natural E6.5, E7.5 and E8.5 and ETiX5, ETiX6 and ETiX8 embryos (n=29 for ETiX5, 10 for ETiX6, 7 for ETiX8, 12 for E5.5, 14 for E6.5, 9 for E8.5). **e.** Annotation of single cell RNA-seq UMAP to show cell types identified in natural and ETiX embryos. **f.** Binning of all sequenced cells in natural and ETiX embryos according to germ layer and embryonic and extraembryonic origin at indicated timepoints. **g.** Density plots highlighting the proportions of tissue types that emerge during natural and ETiX embryo development. **h.** Pearson correlation matrix showing global level of similarity across all identified tissues in natural (rows) in comparison to ETiX embryos (columns).

ETiX embryos faithfully recapitulated the development of natural mouse embryos (Fig. 1b,c, Ext. Fig. 1a-b), including gastrulation. The efficiency of ETiX development from day 4 to day 5 was 21%. Of the structures selected at day 5 for further culture, the efficiency of transitioning from day 5 to 6, day 6 to 7, and day 7 to 8 was greater than 70% (Ext. Fig. 1c). ETiX day 7 embryos displayed a VE-derived yolk sac (Ext. Fig. 1a). Strikingly, ETiX day 7 embryos displayed a clear anterior-posterior axis with bifurcating neural folds extending into a neural tube and culminating in a tail bud, a morphology that resembles the early headfold-stage of an E8.0 natural mouse embryo. Posterior to this, the tail bud joined with allantois tissue which connected to the developing chorion (Fig. 1c). Thus, to our knowledge, these conditions allowed ETiX embryos to develop through and beyond gastrulation to neurulation for the first time.

We also monitored development by examining changes in gene expression. Specifically, we isolated individual ETiX embryos at day 5, day 6, and day 8 and natural embryos at E6.5, E7.5 and E8.5, dissociated them into single cells and performed single cell RNA sequencing (scRNA-seq). UMAP analyses revealed a similar contribution of cells to the developing lineages in natural and ETiX embryos (Fig. 1d). To determine cell types, the cell populations were subclustered using Seurat and subsequently annotated based on published datasets ^11^ (Fig. 1e). We identified 26 tissues based on gene expression patterns, all of which were represented in both natural and ETiX embryo datasets. Some cell types, notably primordial germ cells and neural crest cells, were not detectable by scRNA-seq but were observed by immunofluorescence (see below).

Natural and ETiX embryos displayed a largely conserved distribution of cells between the different germ layers of the epiblast (ectoderm, mesoderm, endoderm) and between embryonic and extraembryonic lineages (epiblast, extraembryonic ectoderm, extraembryonic mesoderm and extraembryonic endoderm) (Fig. 1f). As expected, natural embryos exhibited an increase in cell type complexity over time, corresponding to the formation of differentiated tissues and organs. For instance, cardiomyocytes and neuroectoderm emerged starting from E7.5. Importantly, this increase in cell type complexity and spatiotemporal orders of all the identified populations were comparable in natural and ETiX embryos, indicating that ETiX embryos were on a similar developmental timeline (Fig. 1g). Both systems showed the development of the three germ layers and their derivatives (neuroectoderm, surface ectoderm, gut tube progenitors), the beginning of organogenesis (cardiomyocytes), and the formation of extraembryonic tissues such as amnion and allantois (Fig. 1e and 1g). A Pearson correlation matrix indicated high similarity of gene expression between the tissue clusters of natural and ETiX embryos (Fig. 1h, Ext. Fig.1d). Thus, ETiX embryos recapitulate the generation of the multiple tissues of the neurulating embryo, as evident not only through their morphology, but also their tissue specific gene expression.

## ETiX embryos develop fore- and mid-brain regions

In natural development, the anterior side of the epiblast retains its epithelial character, upregulates expression of the neuroectodermal marker Sox1 from E8.0, and begins the formation of the nervous system. The neuroectodermal lineage gives rise to the forebrain, midbrain, hindbrain and the spinal cord.

To examine neural development, we analysed the expression of well-established neuroectodermal markers by immunofluorescence. Sox1 and Sox2 were expressed in the neuroepithelial cell population, along the entire anteroposterior axis of ETiX day 7 embryos, in a pattern similar to the natural E8.0 embryo (Fig. 2a,b; Ext.Fig. 2a-c). The Sox1-positive neural tube tissue comprised two thirds of the day 7 ETiX embryo and culminated in two neural folds (Ext. Fig. 1a), whereas the Sox1-negative/Brachyury-positive posterior exhibited a tail-bud-like morphology (Fig. 2a,b), reminiscent of natural E8.0 embryos. A Brachyury-positive notochord running below the neural tube was readily apparent in both natural and ETiX embryos (Fig. 2 a,b and Ext. Fig. 2b,c).

**Figure 2:**
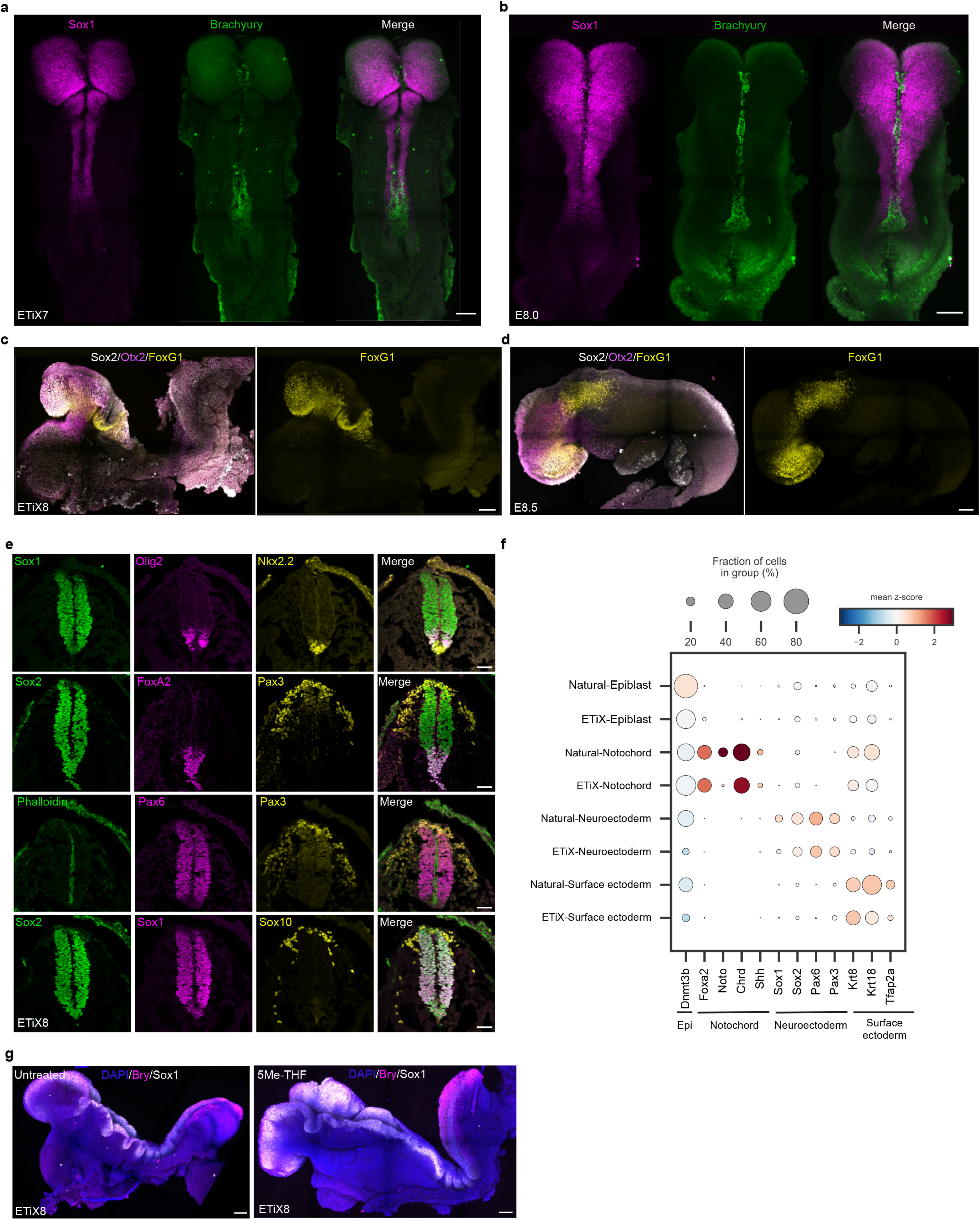
ETiX embryos develop anterior brain structures and a patterned neural tube. Dorsal view of ETiX7 (**a**) and E8.0 natural embryo (**b**) showing formation of Sox1 positive neural folds and the neural tube, which extends from the anterior, with non-overlapping Bry expression, to the Bry-positive posterior (n=11 ETiX7 from 4 independent experiments, n=3 embryos). Lateral view of ETiX8 (**c**) and E8.5 natural embryo (**d**) showing FoxG1 expression in the telencephalon and Otx2 restricted to the forebrain and midbrain and Sox2 marking the whole neural tube (n=4 ETiX8 from 3 independent experiments, n=2 embryos). Scale bar for a,d = 100 μm. **e.** Coronal view of sections of the neural tube showing dorso-ventral patterning at day 8 of ETiX embryo development. Sections are stained to reveal pan-neural markers (Sox1 and Sox2), dorsal markers (Pax6 and Pax3), ventral markers (FoxA2, Olig2 and Nkx2.2) and neural crest markers (Sox10 and Pax3). Scale bar = 50 μm. **f.** Dot plot showing average expression levels and proportion of cells expressing the indicated genes in the selected tissues of both natural and ETiX embryos as indicated based on scRNA-seq data. **g.** ETiX8 embryos following culture in the absence (left) or presence (right) of 5-Methyl-tetrahydrofolate (50 ng/ml) and stained to reveal Sox1, Bry and DNA (n=4 control ETiX, 3 treated ETiX from 2 independent experiments). Scale bar = 100 μm.

The transcription factor Otx2, which is involved in patterning of the midbrain and forebrain^12^ showed restricted expression in the anterior-most third of the head-folds of ETiX day 8 embryos (Fig. 2c,d). This region corresponds to the forebrain and midbrain of the natural mouse embryo at E8.5^13,14^. ETiX day 8 embryos also expressed the transcription factor FoxG1, which plays a vital role in brain development, in the same region as natural E8.5 embryos^15^ (Fig. 2d). The neural tube of ETiX day 8 embryos was closed and showed dorsoventral patterning (Fig. 2e). We detected the distinct neural progenitor domains within the neural tube, delineated by the expression of the markers Pax6, Olig2 and Nkx2.2 ^16–18^ (Fig.2e). Foxa2 was expressed in cells lining the ventral midline of the neural tube, marking a floor plate cell population ^19,20^ (Fig. 2e), whereas Pax3 was expressed in the dorsal neural tube, in the somatic mesoderm and in neural crest cells^21^ (Fig. 2e). Sox10 expression confirmed the identity of neural crest cells^22^, which were displaced from the neural tube, as though undergoing the delamination and migration that occurs during natural brain development^22^ (Fig. 2e).

Next, we examined the scRNA-seq data for the expression of key markers of 1) neuroectoderm, which gives rise to the nervous system, 2) surface ectoderm, which gives rise to most epithelial tissues, 3) notochord, which is important for patterning of the neuroectoderm and neural tube, and 4) epiblast, which is the precursor to all these tissues. In the ETiX embryos and natural notochord, we observed comparable expression of FoxA2, Chordin and Shh (Fig. 2f). Moreover, natural and ETiX embryos expressed similar levels of Sox1, Sox2, Pax6 and Pax3 in the neuroectoderm, and displayed a similar surface ectoderm signature of Keratin gene expression^23^. Thus, tissue-specific gene expression patterns of ETiX embryos strongly resemble those of fertilized natural embryos that were recovered from the mother.

Defects associated with nervous system are the second most frequent developmental abnormality in humans^24–26^, and include brain defects such as anencephaly and spine defects such as spina bifida. We noticed that in certain batches of serum, 75-80% of developing ETiX embryos displayed an abnormal twisting or kinking of the neural tube (Fig. 2g). The vast majority of similar neural tube defects in both mouse and humans can be rescued by supplementing the mother’s diet with folic acid^27^. To test whether folic acid treatment rescues neural tube defects during ETiX embryo development, we cultured the ETiX embryos in the presence of 5-Methyl-THF (50ng/ml), the metabolically active form of folic acid^28^. We observed that 5-Methyl-THF substantially rescued the neural tube defects of ETiX embryos (Fig. 2g), indicating that ETiX embryos can serve as relevant disease models.

## ETiX embryos initiate somitogenesis and heart development

During natural embryogenesis, neuromesodermal progenitors (NMPs) contribute to derivatives of the neural tube and the paraxial mesoderm^29^. To determine if ETiX embryos form NMPs, we performed immunofluorescence. Indeed, we detected expression of the NMP markers Sox2 and Brachyury in a domain spanning the posterior region of the tailbud of ETiX day 8 embryos (Fig. 3a). In contrast, the more anterior regions of ETiX8 embryos expressed Sox2 but not Brachyury, marking the neural lineage, or Brachyury but not Sox2, marking the mesodermal lineage (Fig. 3a). These findings agree with the differentiation trajectory reported for these cells in the natural embryo^30^.

**Figure 3:**
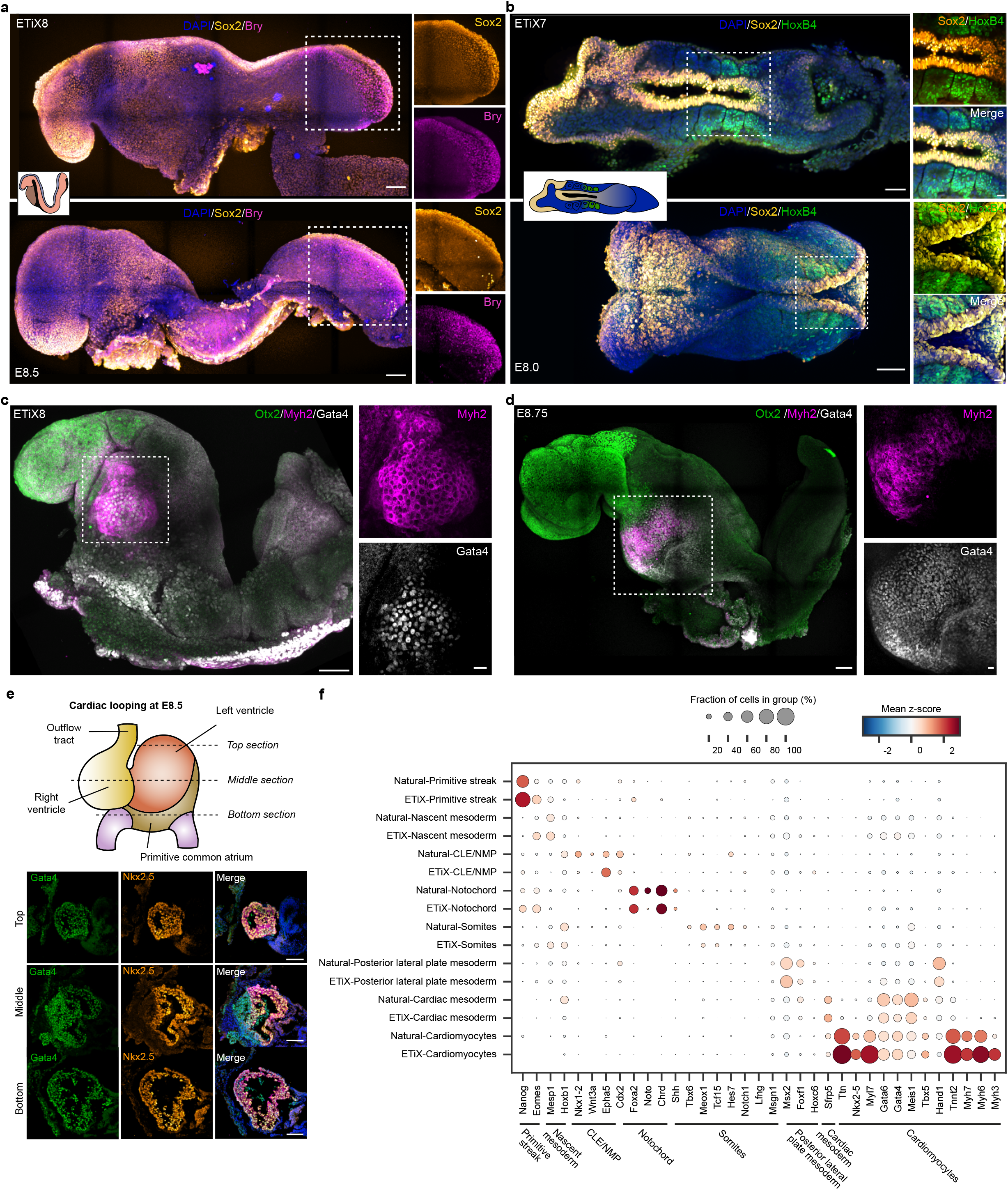
ETiX embryos undertake somitogenesis and heart formation. **a.** (Upper) Lateral view of ETiX8 and (lower) natural E8.5 embryo stained to reveal Sox2, Bry and DNA to highlight formation of neuromesodermal progenitors in the tail bud region (n=5 ETiX8 from 4 independent experiments, n=1 embryo). **b.** (Upper) Dorsal view of ETiX7 and (lower) natural E8.0 embryo stained to reveal Sox2, HoxB4 and DNA to highlight somite formation flanking the neural tube. Enlarged images of somites are shown to the right (n=9 ETiX7 from 4 independent experiments, n=2 embryos). Scale bar for a,b = 100 μm. Scale bar for magnified square (top) = 50 μm, (bottom) = 20 μm **c.** Lateral view of ETiX8 and **(d)** natural E8.75 embryo stained to reveal Otx2, Myh2 and Gata4 to highlight formation of the heart (n=8 ETiX8 from 3 independent experiments, n=1 embryo). Dashed areas are enlarged to the right. Scale bar for c,d = 100 μm. Scale bar for magnified squares = 20 μm. **e.** ETiX8 embryo sectioned coronally. Sections at different depths (top, middle, and bottom) illustrate heart morphogenesis in relation to the natural mouse heart (schematic). Sections are stained to reveal Gata4, Nkx2.5 and Myh2. Scale bar = 100 μm. **f.** Dot plot showing the average expression and proportion of cells expressing the indicated genes in the indicated tissues from natural and ETiX embryos by scRNA-seq.

The paraxial mesoderm in turn gives rise to somites, which are paired blocks of cells that form along the anterior-posterior axis of the developing embryo and are required for segmental formation of skeletal muscle, blood vessels and skin^31^. Pairs of somites expressing the homeobox protein HoxB4 were observed on either side of the Sox1/Sox2-positive neural tube population in ETiX day 7 and natural E8 embryos (Fig. 3b, Extended Fig. 3a-c). These expression patterns mark the key features of somitogenesis, which thus recapitulates the corresponding natural process.

A distinct set of cells destined to form the heart also emerges from the primitive streak at gastrulation. In the natural mouse embryo, this dramatic developmental event takes place around E8.0 and a heartbeat is established as the cardiac mesoderm differentiates into cardiomyocytes. We observed formation of a beating structure below the encephalon region in ETiX day 8 embryos (supplemental Video 1). Myosin heavy chain II (Myh2) and the Gata4 transcription factor are required for cardiac development, and we found that this beating region expressed Myh2 and Gata4 (Fig. 3c, Ext. Fig. 3 d,e) in a similar spatiotemporal profile as the natural embryo (Fig. 3d). Immunostaining of the indicated sections in ETiX day 8 embryos showed a Nkx2.5, Gata4 and Myh2 triple-positive compartment whose architecture resembled the cardiac looping of the developing heart in E8.5 natural embryos (Fig. 3e, Ext. Fig. 3e).

The scRNA-seq data from mesoderm and its derivatives corroborated and extended our findings from immunofluorescence. The gene expression signature leading to somite formation was apparent in the ETiX embryos, although the transcript levels of presomitic identity genes (Tbx6, Hes7 and Msng1), the Notch pathway genes (Notch1, Lfng), and a somite marker (Meox1) were lower than in natural embryos. Both natural and ETiX embryos expressed Gata4 and other important regulators of heart development, including Gata6, Meis1, Tbx5, and Hand1 (Fig. 3f). Similarly, ETiX embryos expressed cardiomyocytes markers such as troponin genes (Ttn and Tnnt2) and myosin genes (Myh7, Myh6, Myl3, Myl7). We infer that mesoderm development in ETiX embryos is remarkably similar to that of natural embryos.

## ETiX embryos initiate gut development

Having observed extensive development and morphogenesis of ectoderm and mesoderm, we next examined development of definitive endoderm, which gives rise to the gut and associated organs. Sox17 is required for gut endoderm development^32^ and whole mount immunofluorescence of ETiX day 8 embryos revealed a Sox17-positive region below the heart, indicative of a primitive gut tube population (Fig.4a,b). By contrast, the heart was Sox17-negative and Gata4-positive, just as in the natural embryo (Fig. 4a,b). Sections of ETiX day 7 embryos revealed an open gut pocket below the neural tube that expressed the transcription factors Sox2 and FoxA2, both critical for gut development (Fig. 4c).

**Figure 4:**
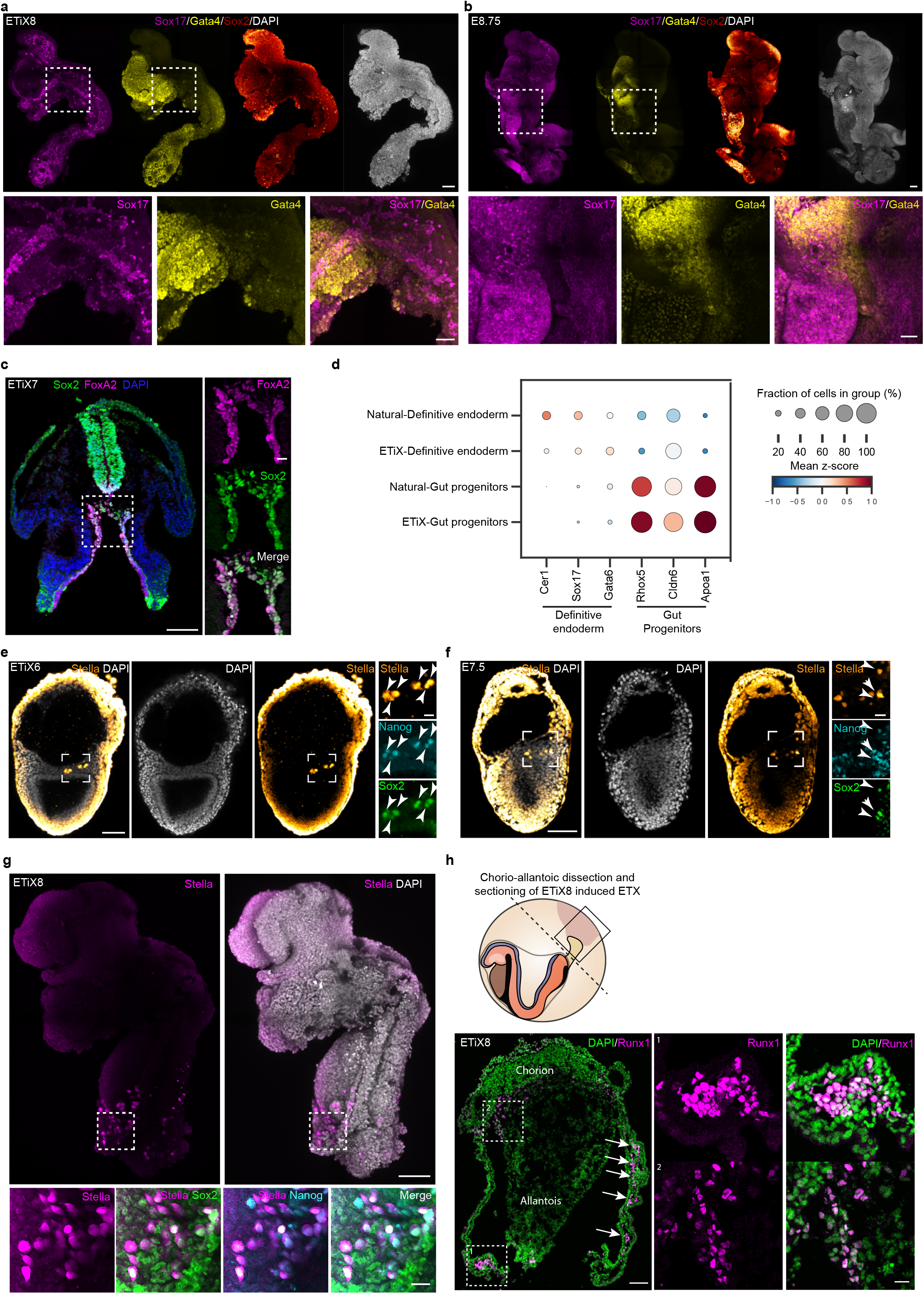
ETiX embryos develop a gut pocket, primordial germ cells and yolk sac blood islands. **a.** ETiX8 and **b.** natural E8.75 embryo stained to reveal Sox17, Gata4 and Sox2 and highlight gut formation in relation to the neural tube (n=6 ETiX8 from 3 independent experiments, n=3 embryos). Boxes indicate magnified regions beneath each panel. Scale bar for a,b = 100 μm. Scale bar for magnified square = 50 μm.**c.** ETiX8 sectioned coronally and stained to reveal Sox2, FoxA2 and DNA to highlight formation of gut pocket in relation to neural tube. Scale bar = 100 μm. Scale bar for magnified square = 20 μm. **d.** Dot plot showing the average expression of, and proportion of cells expressing, indicated genes in selected tissues of natural and ETiX embryos by scRNA-seq. **e.** ETiX6 and **f.** natural E7.5 embryo stained to reveal Stella, Nanog and Sox2 to highlight presence of committed PGCs (n=9 ETiX6 from 2 independent experiments, n=4 embryos). Boxes are shown on the right. **g.** ETiX8 embryo stained to reveal Stella, Nanog and Sox2 to highlight presence of committed PGCs (n=4 ETiX8 from 3 independent experiments). Box is magnified below. **h.** Schematic of yolk sac dissection to isolate the chorioallantoic attachment of ETiX embryos; Sagittal section of chorioallantoic attachment and yolk sac of an ETiX8 embryo stained to reveal Runx1 and DNA. Arrows highlight sites of blood island formation. Boxes are magnified on the right. Scale bar for e-h = 100 μm. Scale bar for magnified square = 20 μm.

The scRNA-seq data from natural and ETiX embryos also revealed gene expression signatures corresponding to definitive endoderm (Cer1/Sox17/Gata6-positive) and VE gut progenitors (Rhox5/Cldn6/Apoa1-positive) (Fig. 4d). Transcripts corresponding to the liver, pancreas, small intestine, or colon were not observed in either ETiX or natural embryos. These data suggest that neither ETiX day 8 nor E8.5 natural embryos have developed beyond an uncommitted endodermal state, consistent with the reported onset of gut tube formation at E8.75^32^.

## ETiX embryos have developing primordial germ cells

Primordial germ cells (PGCs) emerge in the proximo-posterior region of the epiblast, around the same time as Brachyury expression in the E6.5 natural mouse embryo^33,34^. Committed PGCs are characterised by Stella expression^35^, and we detected Stella protein at the ESC-TSC boundary in ETiX day 6 embryos, similarly to the E7.5 natural embryo (Fig. 4e,f). PGCs were also present at later timepoints of ETiX development (Fig. 4g and Ext. Fig 4a,b). During development, PGCs reactivate the pluripotency markers Sox2 and Nanog, and we also observed this to be the case in ETiX embryos (Fig. 4e,f, Ext. Fig. 4a,b) In ETiX embryos at day 7 and day 8, PGCs were localised in proximity to the allantois, in a fashion similar to the natural embryo at E8.0 (Ext. Fig. 4c). Quantification of PGC numbers indicated that ETiX day 8 embryos had 30-100 PGCs, similar to the reported number of PGCs in natural E8.5 embryos^36^ (Ext. Fig. 4d).

## ETiX embryos develop yolk sac blood islands

The extraembryonic portion of the yolk sac in ETiX embryos was attached to a structure resembling the chorion and allantois. The yolk sac supports primitive haematopoiesis in the early embryo, and Runx1 is one of the early markers of haematopoietic progenitors^37^. Strikingly, we observed Runx1-positive blood islands in the mesoderm of the yolk sac and at the base of the allantois of ETiX day 8 embryos (Fig. 4h). Furthermore, the allantois was connected to a chorion-like tissue, which expressed Gata4 and Keratin 18, unlike the yolk sac proper, which expressed only Gata4 (Ext. Fig. 4e).

In conclusion, ETiX embryos have the potential to develop through and substantially beyond gastrulation without the need of providing additional external signalling cues. In contrast to any other stem cell-derived embryo system described to date, ETiX embryos undertake morphogenesis of headfold structures in a manner that closely resembles the natural embryo. We propose that this extended development of ETiX embryos relies on the formation of the anterior signalling centre (namely, the AVE), which protects the anterior embryonic regions from signals that promote posterior development and enables the correct positioning of the primitive streak. Together, these events enable the region immediately anterior to the primitive streak to correctly direct formation of fore- and mid-brain. The natural gastrulation movements of the ETiX embryos enable them to proceed to neurulation with formation of the neural tube, initiation of somitogenesis, and the generation of mesodermal structures including an anatomically accurate beating heart. At present we have not studied development beyond the establishment of the endodermal progenitors for the gut and its associated organs. Notably, in our hands, *ex vivo* culture conditions did not permit the development of natural embryos dissected at E6.5 beyond E8.5 *in vitro*. However, there are no reasons to suspect that, given appropriate culture conditions, ETiX development will not proceed further.

The ability to correct neural tube defects in ETiX embryos highlights their utility as a powerful and tractable mammalian system to discover environmental and genetic factors that predispose offspring to neural tube defects, to dissect the molecular pathways of neural tube formation and closure, and to screen for potential therapeutics. Because ETiX embryos capture extensive development of the epiblast along with the extraembryonic lineages, they provide an unprecedented opportunity to uncover mechanisms of development and disease that involve crosstalk between the next generation and the yolk sac or placenta.

## EXPERIMENTAL MODEL AND SUBJECT DETAILS

### Cell Lines and Culture Conditions

Cell lines used in this study include:

- CAG-GFP/tetO-mCherry mouse ESC (constitutive GFP expression in the membrane; transient mCherry expression upon Dox treatment). The parent CAG-GFP/tetO-mCherry ESC line was derived from an existing mouse line with constitutive CAG-GFP expression and Dox-induced transient mCherry expression. This line was generated by breeding CAG-GFP reporter mice^38^ and tetO-Cherry Histone mice^39^. For the purpose of this study, an independent Dox-inducible Gata4-expressing cassette was introduced into the CAG-GFP/tetO-mCherry ESC line by piggyBac-based transposition, as described below, thus mCherry and Gata4 are regulated by two, independent Dox-responsive promoters.
- CAG-GFP/tetO-mCherry/tetO-Gata4 ESC generated in-house.
- Cerl-GFP mouse ESC (GFP expression under the control of the Cerl-promoter) were derived from a published Cerl-GFP mouse line^40^.
- Cerl-GFP/tetO-Gata4 ESC generated in-house.
- Wildtype TS cells were generated in house from CD1 mice.
- Wildtype CD1 ESC were a kind gift of Dr. Jenny Nichols
- CD1/tetO-Gata4 ESC were generated in-house
- Sox2-Venus/Brachyury-mCherry/Oct4-Venus ESC were a kind gift of Dr. Jesse Veenvliet and Prof. Bernhard G. Hermann.

The sex of the cell lines is not known because we did not genotype them to determine it.

All cell lines were routinely tested every two weeks to ensure that they were not contaminated with mycoplasma.

Mouse ESC were cultured on gelatinised plates at 37°C, 5% CO2, 21% O2 in N2B27 which comprised of 50% Neurobasal-A (Gibco 10888022), 50% DMEM/F-12 (Gibco 21331020), 0.5% N2 (in-house), 1% B27 (Gibco 10889038), 2mM GlutaMAX (Gibco 35050038), 0.1mM 2-mercaptoethanol (Gibco 31350010) and 1% penicillin/streptomycin (Gibco 15140122). N2B27 was supplemented with 3μM CHIR99021 (Cambridge Stem Cell Institute), 1μM PD0325901 (Cambridge Stem Cell Institute) and 10 ng ml^-1^ leukaemia inhibitory factor (Cambridge Stem Cell Institute). Mouse TSC were cultured at 37°C and 5% CO2, in RPMI 1640 (Sigma) with 20% FBS, 2 mM l-glutamine, 0.1 mM 2-ME, 1 mM sodium pyruvate, and 1% penicillin-streptomycin (TSC media), supplemented with 25 ng ml^-1^ FGF4 (R&D Systems 7486-F4-025) and 1 μg ml^-1^ heparin (Sigma-Aldrich H3149-25KU) (TSC/F4H) on mitotically inactivated mouse embryonic fibroblasts (MEFs, Insight Biotechnology, ASF-1201). MEFs were cultured in feeder cell (FC) medium which contained Dulbecco’s modified essential medium (Gibco 41966052), 15% foetal bovine serum (Cambridge Stem Cell Institute), 1mM sodium pyruvate (Gibco 11360039), 2mM GlutaMAX (Gibco 35050038), 1% MEM non-essential amino acids (Gibco 11140035), 0.1mM 2-mercaptoethanol (Gibco 31350010) and 1% penicillin/streptomycin (Gibco 15140122). Passaging of ESC and TSC was performed when they were at 70% confluency as follows: cells were washed once in 1x PBS (Life Technologies 10010056) and trypsinised (Trypsin-EDTA 0.05% Life Technologies 25300054) for 3 minutes at 37°C. The reaction was stopped by adding 2 ml of FC or TSC media respectively, cells were dissociated by pipetting gently 4-5 times and centrifuged for 4 minutes at 200 x g. TSC were then resuspended in TSC/F4H culture media and plated onto MEF-coated plates in 1:20 or 1:10 dilution. ESC were washed once with 1 ml of 1x PBS, centrifuged again, resuspended in N2B27 2iLIF and plated at 1:10 or 1:20 onto gelatine-coated plates.

### Mouse Model

Mice (six-week-old CD-1 males from Charles River and transgenic females bred in house) used in the experiments were kept in animal house, following national and international guidelines. All experiments performed were under the regulation of the Animals (Scientific Procedures) Act 1986 Amendment Regulations 2012 and were reviewed by the University of Cambridge Animal Welfare and Ethical Review Body (AWERB). Experiments were also approved by the Home Office.

## METHOD DETAILS

### Formation of ETiX embryos

To prepare the AggreWell plate (STEMCELL Technologies 34415), 500μl of anti-adherence rinsing solution (STEMCELL Technologies 07010) was added to each well. The plate was then centrifuged at 2,000 x g for 5 minutes and was incubated for 20 minutes at room temperature. Rinsing solution was then aspirated from the well and 1ml of PBS was added to wash each well. 500μl of FC medium was added to each well after aspirating the PBS.

To generate ETiX embryos, doxycycline (1 μg/ml) (Sigma-Aldrich D9891-5G) was added to CAG-GFP tetO-Gata4 ESCs for 6 hours. TSCs were trypsinised and were added to a gelatinised plate to deplete the MEFs for 20 minutes at 37°C. CAG-GFP WT ES cells and CAG-GFP tetO-Gata4 ESCs were subsequently trypsinised. ESCs were washed once with 1x PBS, and trypsinised with 0.05% trypsin-EDTA (ThermoFisher Scientific) for 3 minutes at 37°C. The reaction was stopped by adding 2 ml of FC. Cells were dissociated gently by pipetting for 4-5 times and centrifuged at 200 x g for 4 minutes. The cell pellet was washed once with 1x PBS, centrifuged again and resuspended in 1-2 ml of FC. Cell suspensions with 19,200 TS cells, 6,000 CAG-GFP WT ESC and 6,000 CAG-GFP tetO-Gata4 ESCs were mixed and pelleted by centrifugation. The cell pellet was resuspended in 1ml of FC medium with 7.5nM ROCK inhibitor (Y27632, STEMCELL Technologies 72304). After adding the cell mixture dropwise to the AggreWell, the plate was centrifuged at 100 x g for 3 minutes. On the next day, media change was performed twice by removing 1ml of medium from each well and adding 1ml of fresh FC medium without ROCK inhibitor. On Day 2, media change was performed once to replace 1ml of medium with 1ml of fresh FC medium. On Day 3, 1 ml of medium was removed from each well and 1.5ml of IVC1 was added, after equilibrating for 20 minutes in the incubator. IVC1^41^ is made of advanced DMEM/F12 (Gibco, 21331-020) supplemented with 20% (v/v) FBS, 2 mM GlutaMax, 1% v/v penicillin–streptomycin, 1X ITS-X Thermo Fisher Scientific, 51500-056), 8 nM β-estradiol, 200 ng/ml progesterone and 25 μM N-acetyl-L-cysteine. On Day 4, ETiX embryos in the AggreWell were transferred to CELLSTAR 6 well multiwell plate for suspension culture (Greiner Bio-One 657185) with 5ml of IVC1 (with FBS at 30% v/v) per well. On Day 5 IVC1 was replaced with *ex utero* culture medium (EUCM). EUCM comprises 25% DMEM (GIBCO 11880) supplemented with 1× Glutamax (GIBCO, 35050061), 100 units/ml penicillin and 100 μg/ml streptomycin and 11 mM HEPES (GIBCO 15630056), plus 50% rat serum (rat whole embryo culture serum, Charles River) and 25% human chord serum obtained from the Cambridge Blood Biobank. Human and rat serum were heat-inactivated for 35 minutes (from frozen) at 56 C and sterilised by filtration^10^. Each ETiX was transferred to a single well of 24-well, non-adherent dish (Greiner 662102) with 250 uL of EUCM. On day 6 each ETiX embryos was fed with an additional 250 uL of EUCM. On day 7, EUCM was supplemented with 3.0 mg/ml of D-Glucose (Sigma G8644) and the samples were moved to a rotating bottle culture chamber apparatus^10^. Each rotating bottle contained 2 ml of EUCM and 3 ETiX embryos. On day 8 EUCM was supplemented with 3.5 mg/ml of D-Glucose; in each rotating bottle 2 ETiX were cultured with 3 ml of EUCM.

### Recovery of Mouse Embryos

Natural mating was performed with six-week-old transgenic females and CD-1 males. Mouse embryos were recovered at embryonic days E5.5, E6.5 and E7.5 by dissecting from the deciduae in M2 medium (Sigma M7167). Embryos at E6.5 were cultured in EUCM in stationary conditions until E8.5 as reported^10^ in the same way as the ETiX. At E8.5, embryos were moved to a rotating bottle in EUCM supplemented with 3.0 mg/ml of D-glucose. Each bottle contained 2 ml of EUCM medium and 3 embryos.

### Plasmids and Transfection

Gata4 cDNA was PCR-amplified from pSAM2-mCherry-Gata4 using the Gata4/AttB primers (see table). The primers were designed as outlined in the Gateway cloning manual. Because the plasmid already contained attB sites, there was no need to incorporate parts of the Gata4 open reading frame in the primer design. pSAM2-mCherry-Gata4 was a gift from Timothy Kamp (Addgene plasmid # 72690; http://n2t.net/addgene:72690; RRID:Addgene_72690,)^42^. It was subsequently cloned into PB-tetO-hygromycin by Gateway technology (Thermo Fisher Scientific), according to the manufacturer’s instructions. Clones were verified by sequencing. Transformations were performed using 5α-competent *E.coli* (New England Biolabs C2987I). To generate ES cells with Doxycycline inducible Gata4, PB-tetO-hygro-Gata4, pBAse and rtTA-zeocyin (0.25 μg/each/reaction) were transfected into 12,000 CAG-GFP ES cells using Lipofectamine 3000 Transfection Reagent (Invitrogen L3000001), followed by antibiotics selection for 7 days with hygromycin (1:250; Gibco 10687010) and zeocyin (1:1000; InvivoGen ant-zn-1). The PB-tetO-hygro, pBAse and rtTA-zeocyin were generously gifted by Dr. Jose Silva from the Stem Cell Institute (Cambridge, UK).

### Immunofluorescence

ETiX embryos and natural mouse embryos were fixed with 4% paraformaldehyde at room temperature for 20 minutes and washed with PBST (PBS with 0.1% Tween 20) for three times. They were then permeabilised in permeabilization buffer (0.1 M glycine and 0.3% Triton X-100 in PBST). For embryos up to E7.5 and ETiX embryos up to day 6, permeabilization was performed for 30 minutes at room temperature, followed by three washes with PBST of five minutes each. Older natural and ETiX embryos were permeabilised for 35 minutes. Samples were incubated with primary antibodies diluted in blocking buffer (10% FBS and 0.1% Tween 20 in PBS) at 4°C overnight. After washing with PBST for three times, they were incubated with secondary antibodies and DAPI at 4°C overnight or for 2 hours at room temperature followed by another three washes with PBST before imaging.

#### Cryosectioning and slide immunofluorescence

Natural and ETiX-embryo samples after fixation were cryoprotected in 30% sucrose/PBS (w/v) overnight at 4C or until the sample sank to the bottom of the tube. Samples were then transferred to a cryomold. The cryomold was carefully filled with OCT compound (Agar Scientific) and frozen on a metal block cooled down in dry ice. The samples were cut at a thickness of 12 um on a cryostat, collected on lysine coated slides and stored at −80 until ready for immunofluorescence. Immunofluorescence was performed as follows: the slides were washed in the PBS to remove the OCT for 5-10 minutes and then briefly allowed to air dry for an additional 5-10 minutes. Samples were permeabilised for 10 minutes at room temperature with permeabilization buffer (0.1 M glycine and 0.3% Triton X-100 in PBST) and then blocked for 1 hour at room temperature with blocking buffer (10% FBS and 0.1% Tween 20 in PBS). After permeabilization and blocking the samples were washed for 3 times in PBS for 5 minutes at room temperature. Primary antibodies were diluted in blocking buffer and incubated with the sample overnight at 4C. Following 3 washes of 5 minutes each in PBS, the sample was incubated for 2 hours at room temperature with secondary antibodies diluted in blocking buffer. After three final washes, slides were mounted with homemade mounting media, sealed with nail polish, and allowed to airdry overnight in the dark prior to imaging.

### scRNA-seq Sample Preparation and Dissociation

After recovery, natural embryos and ETiX embryos were cut to pieces, transferred to a Falcon tube, centrifuged, washed in PBS and incubated in Tryple Express (Gibco™ 12604013) for 15 minutes, with vigorous pipetting for 20 times every 5 minutes to dissociate to single cells. If there were clumps left, the incubation was extended for an additional 5 minutes and the sample was pipetted further. Samples were filtered with a 40 um filter to remove large clumps, centrifuged at 200 x g for 5 minutes and resuspended in PBST (PBS with 0.02% Tween20). The suspension was quantified with Trypan Blue (Sigma T8154) in a 1:1 ratio to determine the proportion of live and dead cells and then processed for encapsulation (see below).

### scRNA-seq Library Preparation and Sequencing

Libraries were prepared according the inDrops v3^43^ workflow^44^ with v3 barcoding scheme^45^. Briefly, polyacrylamide beads were generated and barcoded to obtain a diversity of 147,456 barcodes. Single-cell suspensions were diluted to a concentration of 100,000 cells per ml and co-encapsulated with the barcoded beads and reverse transcriptase and lysis mix. Fractions of ~1,000 cells were collected in 1.5 ml Eppendorf tubes pre-filled with 200 μl of mineral oil, subjected to UV photocleavage and incubated at 50C for 2 hours and 70C for 20 minutes. The droplets were then de-emulsified and further amplified using second-strand synthesis and *in vitro* transcription. The libraries were then fragmented and reverse transcribed. The final libraries were amplified using a unique 8bp index using limited-cycle PCR and quantified using a Qubit High sensitivity kit (Invitrogen) and Bioanalyzer High sensitivity DNA kit (Agilent). Libraries were pooled at equi-molar ratios and purified using a 1.5x volumetric ratio of AmpureXP beads. The libraries were sequenced on a Nextseq 75 cycle 400M read High Output kit with 5% PhiX spike-in as an internal control. The read cycle distribution was the following: Read1 61 cycles; Index1 8 cycles; Index2 8 cycles; Read2 14 cycles.

### scRNA-seq Bioinformatic Analysis

The BCL files were converted to Fastq files using Illumina’s bcl2fastq software. The sequenced libraries were quality-inspected using the FastQC^46^ tool and de-multiplexed using the Pheniqs tool from biosails. The fastq files were further filtered, mapped to a mouse GRCm38.99 reference genome with GRCm38.99 gtf annotation and deduplicated using the zUMIs pipeline^47^. The count matrices with exonic and intronic counts were then used as an input for downstream analysis using Seurat^48^. Cells were filtered based on the number of genes detected (between 700 and 4,000), UMIs detected (lower than 7,500), percentage of UMI counts mapping to mitochondrial genes (between 1 and 15%) and doublet scores computed using Scrublet^49^ (lower than 0.3), which yielded a total of 26,748 cells overall. The natural and synthetic embryo datasets were integrated in Seurat and shared embeddings were corrected for batch effect (both systems and timepoints collected) using Harmony^50^. Louvain clustering was performed on the shared embeddings and markers, computed using the FindAllMarkers function from Seurat, were used to annotate cell types. Pearson correlation coefficients between cell types for each system, single-cell velocity profiles and latent times were computed using the Scanpy^51^ and scVelo^52^ tools.

### Inclusion Criteria of ETiX embryos

All ETiX embryos were collected from AggreWell for analysis at 4 days of development and analysed under a stereo microscope. In all instances, we selected ETiX embryos with cylindrical morphology and two clearly defined cellular compartments surrounded by an outer cell layer. We expect the ESC compartment to be epithelialized with a lumen. The TSC compartment is more variable in appearance and therefore, even though one would also want an epithelial-looking TSC compartment similar to the extra-embryonic ectoderm of natural embryos, we select a wider range of appearances for the TSC compartment. After this initial selection, structures containing the appropriate fluorophores are quickly checked under a microscope to ensure the presence of an epithelialized CAG-GFP-positive ESC compartment. When ETiX were generated by using wild-type stem cell lines, the selection was based on morphology alone. ETiX embryos with the correct body plan of ESC and TSC compartments surrounded by a visceral endoderm-like layer are then transferred to equilibrated media to continue their culture. When selecting at D5, however, we included additional criteria: i) we expect the lumen of the ESC and TSC compartment to be merged; ii) ideally we can observe the beginning of gastrulation on one side of the ETiX embryo; iii) we expected the AVE to have migrated to the ESC-TSC boundary and be opposite to the forming streak; iv) ETiX with the AVE stuck at the tip of the structure or not at the boundary were excluded; v) when using the Triple reporter cells, we ensured that one side showed higher Sox2-Venus expression.

### Image Acquisition, Processing and Analysis

Images were acquired using Leica SP5 and SP8 confocal microscopes (Leica Microsystems) with 40× oil objective and 25× water objective, respectively. A 405 nm diode laser (DAPI), a 488 nm argon laser (Alexa Fluor 488), a 543 nm HeNe laser (Alexa Fluor 568) and a 633 nm HeNe laser (Alexa Fluor 647) were used to excite the fluorophores. Images were taken with a z-step of 1.2-5μm. Fiji^53^ and NDSAFIR 3.0^54^, Adobe Illustrator 26.0.1 and the Smart Denoise plugin were used to process and analyse the images.

## Acknowledgements

We would like to thank all the members of MZG lab for their helpful comments throughout this project. This project has been made possible through the following grants to MZG: NIH Pioneer Award (DP1 HD104575-01), European Research Council (669198), the Wellcome Trust (207415/Z/17/Z), Open Philanthropy/Silicon Valley Community Foundation and Weston Havens Foundation and the Centre for Trophoblast Research. FH was supported by the ERC (69566) and the Wellcome Trust (); and JdJ thanks the BBSRC DTP for a studentship. CEH was supported by the Centre for Trophoblast Research, and the Leventis Foundation.

## Contributions

GA and CEH designed and carried out the experiments and data analysis. JdJ performed the library preparation for single cell RNA-sequencing and bioinformatics analysis. FH supervised the single cell RNA-sequencing analysis. MZG, GA, CEH wrote the manuscript with DG. MZG conceived and supervised the study.

## RESOURCE AVAILABILITY

### Lead Contact

Further information and requests for resources and reagents should be directed to Magdalena Zernicka-Goetz.

### Materials Availability

All unique/stable reagents generated in this study are available from the Lead Contact with a completed Materials Transfer Agreement.

### Code Availability

Any custom code generated in this study is available upon request.

#### Table of Antibodies

| NAME | SOURCE | IDENTIFIER |
| --- | --- | --- |
| Rabbit polyclonal anti-GATA-4 | Santa Cruz Biotechnology | Cat# sc-9053, RRID:AB_2247396 |
| Goat polyclonal anti-GATA-6 | R&D Systems | Cat# AF1700, RRID:AB_2108901 |
| Goat polyclonal anti-Otx2 | R&D Systems | Cat# AF1979, RRID:AB_2157172 |
| Rabbit monoclonal anti-RUNX1 / AML1 | Abcam | Cat# ab92336, RRID:AB_2049267 |
| Goat polyclonal anti-Sox17 | R&D Systems | Cat# AF1924, RRID:AB_355060 |
| Rabbit monoclonal anti-HoxB4 | Abcam | Cat# ab133521 |
| Rabbit polyclonal anti-Sox1 | Cell Signaling Technology | Cat# 4194 RRID:AB_1904140 |
| Goat polyclonal anti-Sox1 | R&D Systems | Cat# AF3369 RRID:AB_2239879 |
| Goat polyclonal anti-Brachyury | R&D Systems | Cat# AF2085, RRID:AB_2200235 |
| Mouse monoclonal anti-Sox2 | Santa Cruz Biotechnology | Cat# sc-365823 RRID:AB_10842165 |
| Rat monoclonal anti-Sox2 | Thermo Fisher Scientific | Cat# 14-9811-82 RRID:AB_11219471 |
| Rabbit monoclonal anti-FoxG1 | Abcam | Cat# ab196868 RRID:AB_2892604 |
| Rabbit monoclonal anti-FoxA2 / HNF3 | Cell Signaling Technology | Cat# 8186, RRID:AB_10891055 |
| Mouse monoclonal anti-Myh2 | R&D Systems | Cat# MAB4470 |
|  |  | RRID:AB_1293549 |
| Goat polyclonal anti-Nkx2.5 | R&D Systems | Cat# AF2444<br>RRID:AB_355269 |
| Rabbit polyclonal anti-Nanog | Abcam | Cat# ab80892<br>RRID:AB_2150114 |
| Mouse monoclonal anti-Pax3 | DSHB | Cat# AB_528426<br>RRID:AB_528426 |
| Rabbit polyclonal anti-Pax6 | BioLegend | Cat# PRB-278P<br>RRID:AB_291612 |
| Rabbit anti-Olig2 | IBL America | Cat# 18953<br>RRID:AB_1630817 |
| Mouse monoclonal anti Nkx2.2 | DSHB | Cat# 74.5A5<br>RRID:AB_531794 |
| Mouse monoclonal anti-Sox10 | Santa Cruz<br>Biotechnology | Cat# sc-365692<br>RRID:AB_10844002 |
| Goat polyclonal anti-Stella | R&D Systems | Cat# AF2566<br>RRID:AB_2094147 |
| Rabbit polyclonal anti-Keratin18 | Sigma-Aldrich | Cat# SAB4501665<br>RRID:AB_10746153 |
| Alexa Fluor® 488 Phalloidin | Thermo Fisher<br>Scientific | Cat# A12379 |
| Alexa Fluor® 594 Phalloidin | Thermo Fisher<br>Scientific | Cat# A12381 |
| Alexa Fluor® 647 Phalloidin | Thermo Fisher<br>Scientific | Cat# A22287 |
| Donkey anti-Mouse IgG (H+L) Highly Cross-Adsorbed Secondary Antibody, Alexa Fluor 488 | Thermo Fisher<br>Scientific | Cat# A-21202,<br>RRID:AB_141607 |
| Donkey anti-Rabbit IgG (H+L) Highly Cross-Adsorbed Secondary Antibody, Alexa Fluor 488 | Thermo Fisher<br>Scientific | Cat# A-21206,<br>RRID:AB_2535792 |
| Donkey anti-Goat IgG (H+L) Cross-Adsorbed Secondary Antibody, Alexa Fluor 488 | Thermo Fisher<br>Scientific | Cat# A-11055,<br>RRID:AB_2534102 |
| Donkey anti-Mouse IgG (H+L) Highly Cross-Adsorbed Secondary Antibody, Alexa Fluor 568 | Thermo Fisher<br>Scientific | Cat# A10037,<br>RRID:AB_2534013 |
| Donkey anti-Rabbit IgG (H+L) Highly Cross-Adsorbed Secondary Antibody, Alexa Fluor 568 | Thermo Fisher<br>Scientific | Cat# A10042,<br>RRID:AB_2534017 |
| Donkey anti-Goat IgG (H+L) Cross-Adsorbed Secondary Antibody, Alexa Fluor 568 | Thermo Fisher<br>Scientific | Cat# A-11057,<br>RRID:AB_2534104 |
| Donkey anti-Mouse IgG (H+L) Highly Cross-Adsorbed Secondary Antibody, Alexa Fluor 647 | Thermo Fisher<br>Scientific | Cat# A-31571,<br>RRID:AB_162542 |
| Donkey anti-Rabbit IgG (H+L) Highly Cross-Adsorbed Secondary Antibody, Alexa Fluor 647 | Thermo Fisher<br>Scientific | Cat# A-31573,<br>RRID:AB_2536183 |
| Donkey anti-Goat IgG (H+L) Cross-Adsorbed Secondary Antibody, Alexa Fluor 647 | Thermo Fisher<br>Scientific | Cat# A-21447,<br>RRID:AB_2535864 |
| Donkey Anti-Rat IgG H&L (Alexa Fluor® 647) preadsorbed antibody | Abcam | Cat# ab150155,<br>RRID:AB_2813835 |

**Extended Figure 1:**
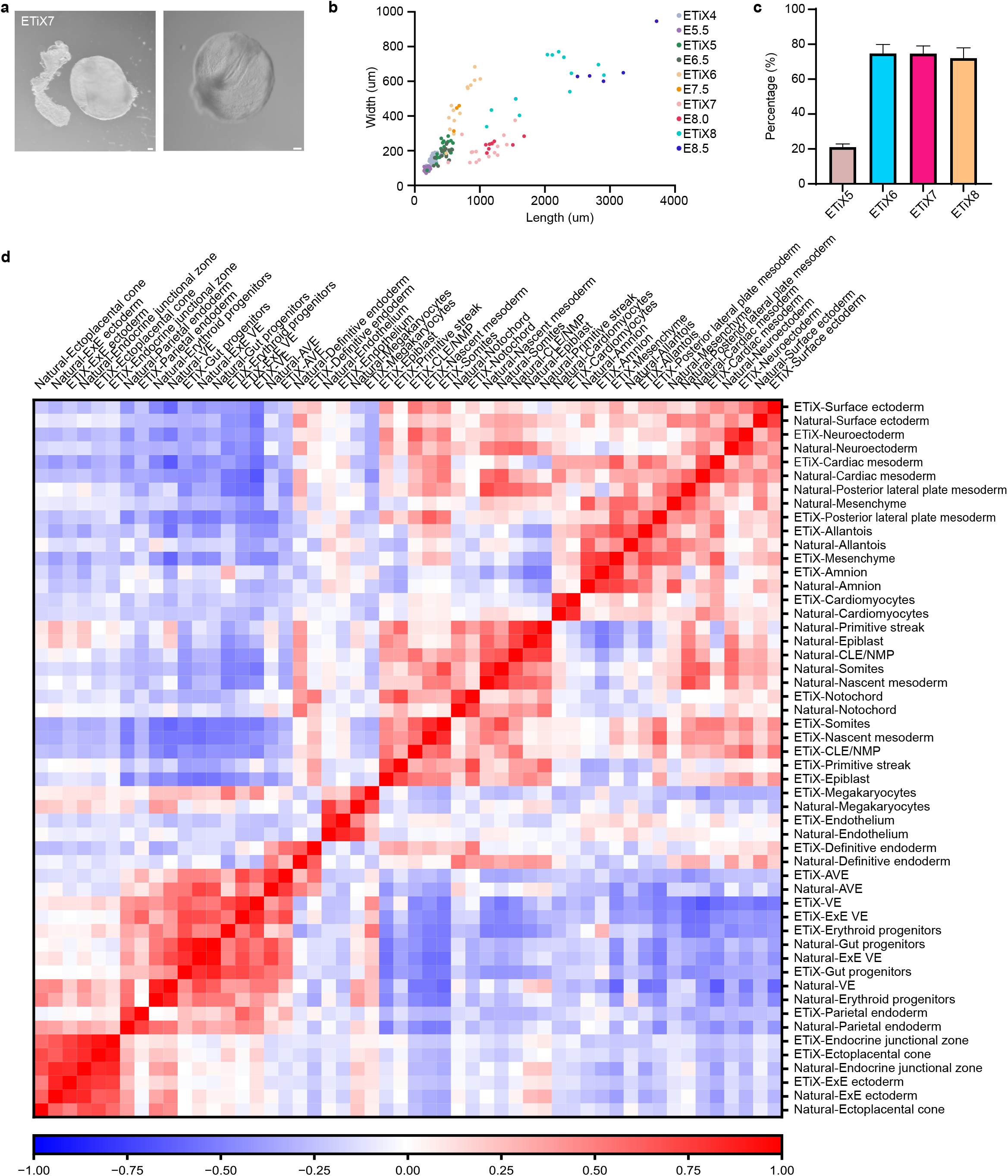
ETiX embryos develop to comparable sizes and gene expression as natural embryos with reproducible efficiency. Brightfield images of ETiX embryos at day 7 prior to dissection to highlight the presence of a yolk sac. Scale bar = 100 μm. **b.** Quantification of ETiX and natural embryo dimensions at comparable developmental timepoints (n=42 for ETiX4, 24 for ETiX5, 14 for ETiX6, 18 for ETiX7, 12 for ETiX8, 32 for E5.5, 18 for E6.5, 3 for E7.5, 8 for E8.0, 5 for E8.5, from 30 independent experiments). **c.** Quantification of ETiX embryo formation efficiency from day 5 to day 8 (n=1197 ETiX4, 237 for ETiX5, 170 for ETiX6, 100 for ETiX7, 40 for ETiX8, from 17 independent experiments). Plots show the mean (+SEM). **d.** Pearson correlation matrix showing the global level of similarity across all comparisons of tissue types in natural and ETiX embryos.

**Extended Figure 2.**
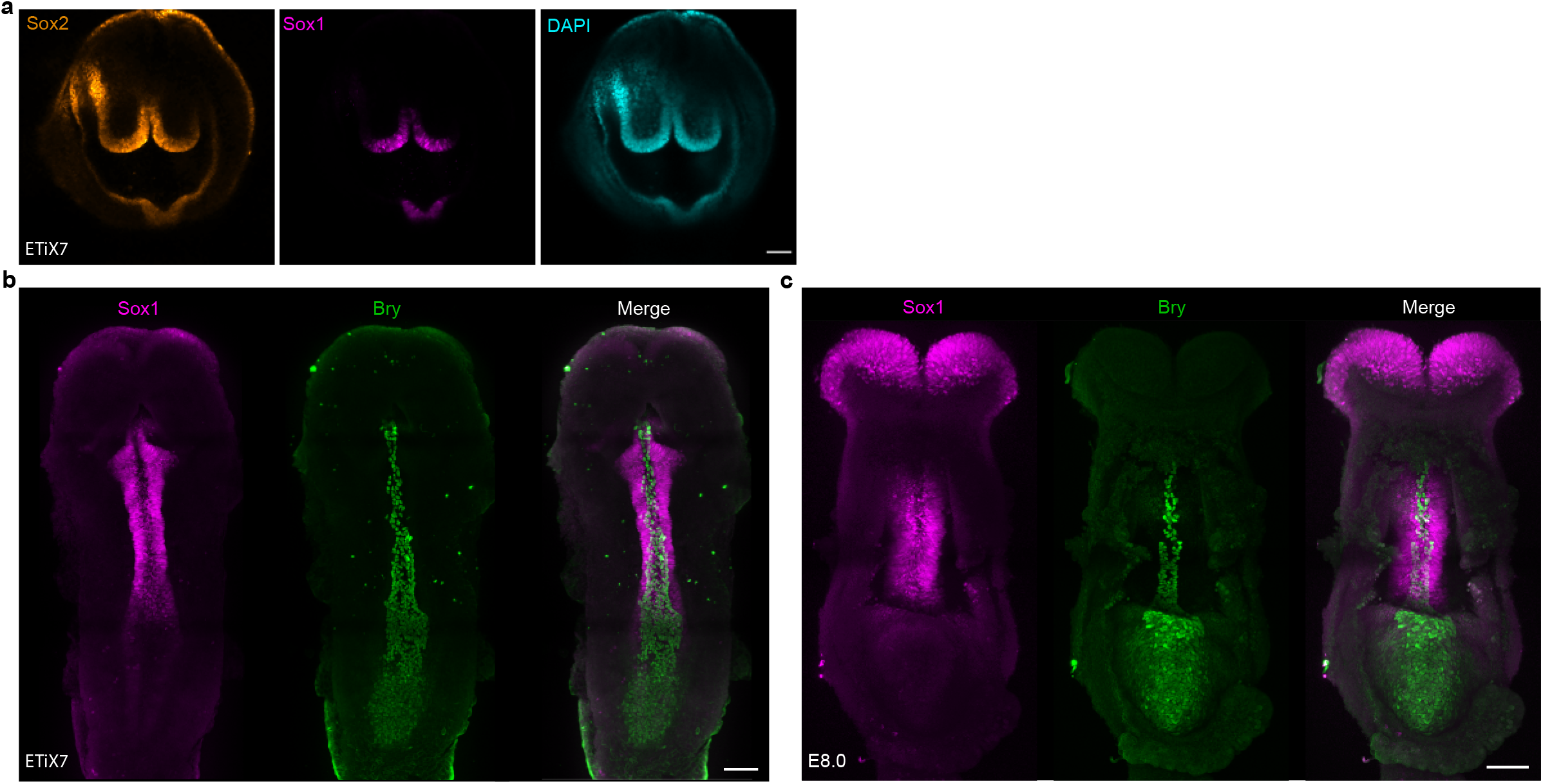
ETiX embryos show neural folds and a developing tail bud. **a.** ETiX7 embryo stained to reveal Sox2, Sox1 and DNA highlighting formation of the rostral neural folds (n=11 ETiX7 from 4 independent experiments, n=3 natural embryos). Ventral views of ETiX7 (**b**) and natural (**c**) E8.0 embryos shown in Fig. 2a, 2b showing formation of Sox1 positive neural folds and Bry-positive notochord and tail bud (n=11 ETiX7 from 4 independent experiments, n=3 embryos). Scale bar = 100 μm.

**Extended Figure 3:**
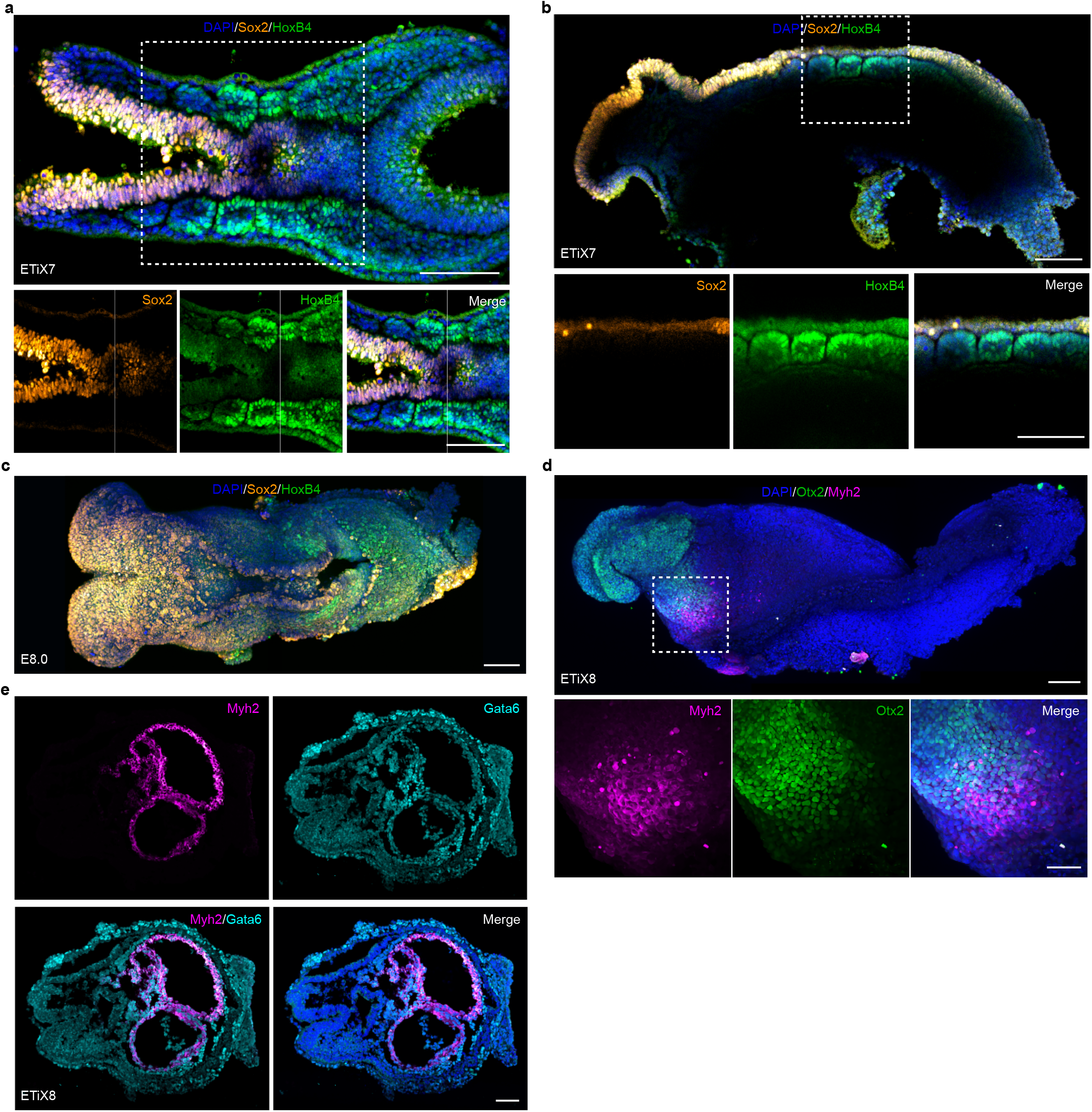
Development of ETiX embryos mesoderm into somites and cardiac tissue. **a.** Dorsal and **b.** lateral view of ETiX7 embryo stained to reveal Sox2, HoxB4 and DNA to highlight somite formation flanking the neural tube (n=9 ETiX7 from 4 independent experiments). **c.** Dorsal view of natural E8.0 embryo stained to reveal Sox2, HoxB4 and DNA to highlight somite formation flanking the neural tube (n=2 embryos). **d.** Lateral view of ETiX8 embryo stained to reveal Otx2, Myh2 and DNA to highlight heart formation (n=8 ETiX8 from 3 independent experiments). Scale bar for a-d = 100 μm. Scale bar for magnified square in d = 50 μm. **e.** ETiX8 embryo sectioned coronally and stained to reveal Gata6 and Myh2 to highlight heart morphogenesis. Scale bar = 200 μm.

**Extended Figure 4:**
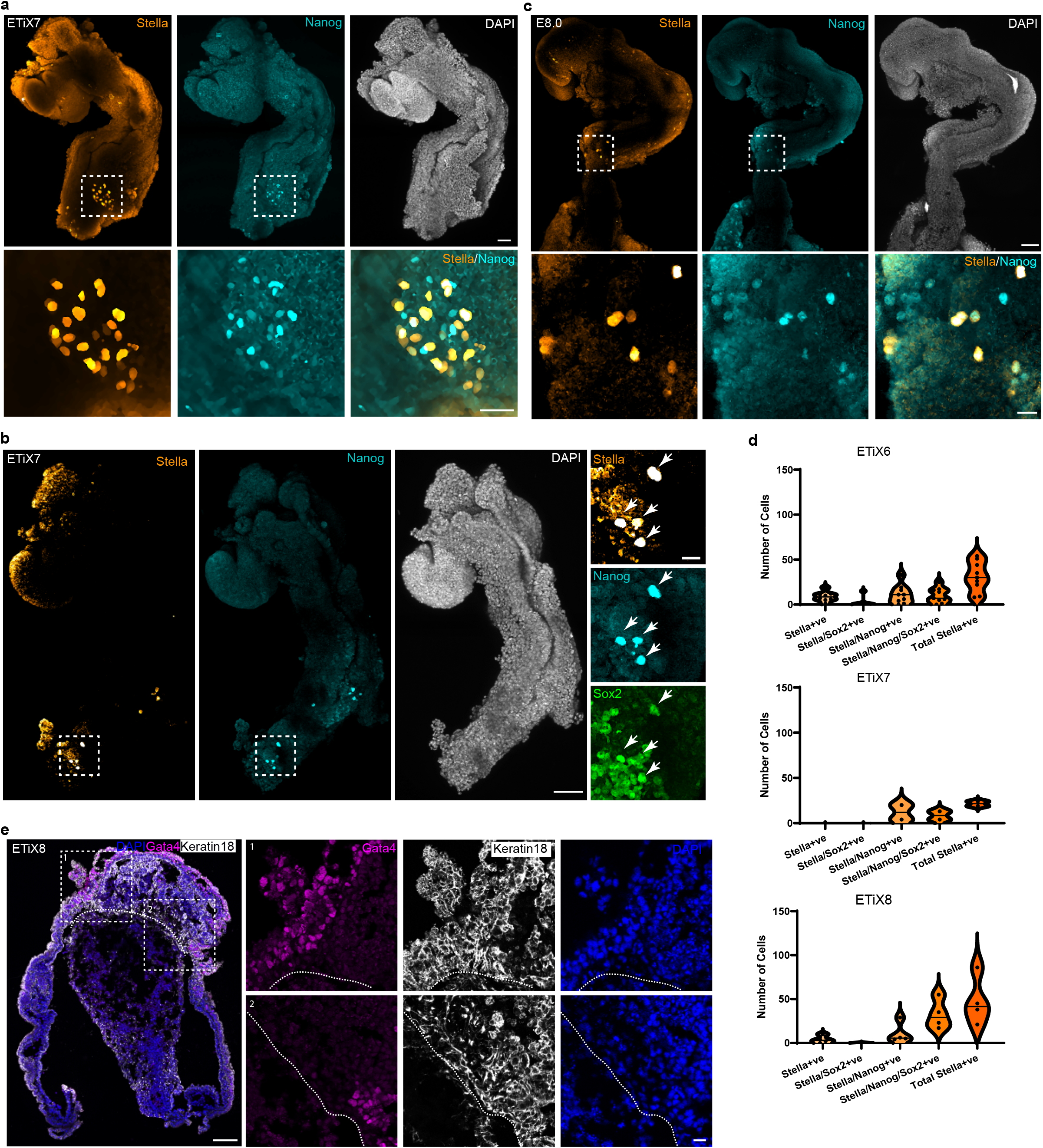
Primordial germ cell numbers in ETiX embryos and development of extraembryonic tissues. **a, b** ETiX-embryo at day 7 of development and (**c)** natural embryo at E8.0 of development analysed for Stella, Nanog and Sox2 to highlight presence of committed PGCs (n=2 structures from 2 independent experiments, n=2 embryos). Boxes are magnified below (a,c) and on the right (b). **d.** Quantification of PGC formation at different stages of ETiX embryos development (n=9 for ETiX6, 2 for ETiX7, 4 for ETiX8). PGCs were scored for Stella expression and co-expression of Nanog and Sox2. Data are presented as violin plots with median and quartiles. **e.** Sagittal section of chorioallantoic attachment and yolk sac for an ETiX8 embryo stained for Keratin18, Gata4 and DNA to visualise chorion and yolk sac. Scale bar for a-c and e = 100 μm. Scale bar for magnified square = 20 μm.

**Video 1: Live imaging of ETiX8 embryo showing beating heart** n=11 structures from 4 independent experiments.

**Video 2: Live imaging of ETiX8 embryo showing beating heart** n=11 structures from 4 independent experiments.

## References

1. van den Brink, S. C. et al. Symmetry breaking, germ layer specification and axial organisation in aggregates of mouse embryonic stem cells. Development 141, 4231–4242 (2014).

2. Beccari, L. et al. Multi-axial self-organization properties of mouse embryonic stem cells into gastruloids. Nature 562, 272–276 (2018).

3. Turner, D. A. et al. Anteroposterior polarity and elongation in the absence of extra-embryonic tissues and of spatially localised signalling in gastruloids: mammalian embryonic organoids. Development 144, 3894–3906 (2017).

4. van den Brink, S. C. et al. Single-cell and spatial transcriptomics reveal somitogenesis in gastruloids. Nature 582, 405–409 (2020).

5. Veenvliet, J. et al. Mouse embryonic stem cells self-organize into trunk-like structures with neural tube and somites. Science 370, eaba4937 (2020).

6. Girgin, M. U. et al. Bioengineered embryoids mimic post-implantation development in vitro. Nature Communications 12, 5140 (2021).

7. Xu, P.-F. et al. Construction of a mammalian embryo model from stem cells organized by a morphogen signalling centre. Nature Communications 12, 3277 (2021).

8. Sozen, B. et al. Self-assembly of embryonic and two extra-embryonic stem cell types into gastrulating embryo-like structures. Nature Cell Biology 20, 979–989 (2018).

9. Amadei, G. et al. Inducible Stem-Cell-Derived Embryos Capture Mouse Morphogenetic Events In Vitro. Developmental Cell 56, 366–382.e9 (2021).

10. Aguilera-Castrejon, A. et al. Ex utero mouse embryogenesis from pre-gastrulation to late organogenesis. Nature 593, 119–124 (2021).

11. Pijuan-Sala, B. et al. A single-cell molecular map of mouse gastrulation and early organogenesis. Nature 566, 490–495 (2019).

12. Acampora, D. et al. OTD/OTX2 functional equivalence depends on 5’ and 3’ UTR-mediated control of Otx2 mRNA for nucleo-cytoplasmic export and epiblast-restricted translation. Development 128, 4801–4813 (2001).

13. Ang, S.-L., Conlon, R. A., Jin, O. & Rossant, J. Positive and negative signals from mesoderm regulate the expression of mouse Otx2 in ectoderm explants. Development 120, 2979–2989 (1994).

14. Pevny, L. H., Sockanathan, S., Placzek, M. & Lovell-Badge, R. A role for Sox1 in neural determination. Development 1967–1978 (1998) doi:https://doi.org/10.1016/j.isci.2020.101475.

15. Hettige, N. C. & Ernst, C. FOXG1 Dose in Brain Development. Frontiers in Pediatrics vol. 7 (2019).

16. Ericson, J. et al. Pax6 Controls Progenitor Cell Identity and Neuronal Fate in Response to Graded Shh Signaling concentrations of Shh are required for the induction of V1 and V2 interneurons and for MNs, with the requisite. Cell vol. 90 (1997).

17. Briscoe, J. et al. Homeobox gene Nkx2.2 and specification of neuronal identity by graded Sonic hedgehog signalling. Nature 389, 622–627 (1999).

18. Novitch, B. G., Chen, A. I. & Jessell, T. M. Coordinate Regulation of Motor Neuron Subtype Identity and Pan-Neuronal Properties by the bHLH Repressor Olig2. Neuron vol. 31 (2001).

19. Ribes, V. et al. Distinct Sonic Hedgehog signaling dynamics specify floor plate and ventral neuronal progenitors in the vertebrate neural tube. Genes and Development 24, 1186–1200 (2010).

20. Sasaki, H. & Hogan, B. L. M. HNF-3/l as a Regulator of Floor Plate Development. Cell vol. 76 (1994).

21. Serbedzija, G. N. & Mcmahon, A. P. Analysis of Neural Crest Cell Migration in Splotch Mice Using a Neural Crest-Specific LacZ Reporter. DEVELOPMENTAL BIOLOGY vol. 185 (1997).

22. Southard-Smith, E. M., Kos, L. & Pavan, W. J. Sox10 mutation disrupts neural crest development in Dom Hirschsprung mouse model. http://www.nature.com/naturegenetics (1998).

23. Tadeu, A. M. B. & Horsley, V. Notch signaling represses p63 expression in the developing surface ectoderm. Development 140, 3777–3786 (2013).

24. Burren, K. A. et al. Gene-environment interactions in the causation of neural tube defects: folate deficiency increases susceptibility conferred by loss of Pax3 function. Human molecular genetics 17, 3675–3685 (2008).

25. Copp, A. J., Greene, N. D. E. & Murdoch, J. N. The genetic basis of mammalian neurulation. Nature Reviews Genetics 4, 784–793 (2003).

26. Copp, A. J., Stanier, P. & Greene, N. D. E. Neural tube defects: recent advances, unsolved questions, and controversies. The Lancet Neurology 12, 799–810 (2013).

27. DiCicco-Bloom, E. et al. The Developmental Neurobiology of Autism Spectrum Disorder. The Journal of Neuroscience 26, 6897 (2006).

28. Wang, X. & Fenech, M. A comparison of folic acid and 5-methyltetrahydrofolate for prevention of DNA damage and cell death in human lymphocytes in vitro. Mutagenesis vol. 18 https://academic.oup.com/mutage/article/18/1/81/1118046 (2003).

29. Tzouanacou, E., Wegener, A., Wymeersch, F. J., Wilson, V. & Nicolas, J. F. Redefining the Progression of Lineage Segregations during Mammalian Embryogenesis by Clonal Analysis. Developmental Cell 17, 365–376 (2009).

30. Koch, F. et al. Antagonistic Activities of Sox2 and Brachyury Control the Fate Choice of Neuro-Mesodermal Progenitors. Developmental Cell 42, 514–526.e7 (2017).

31. Pourquié, O. 3 Segmentation of the Paraxial Mesoderm and Vertebrate Somitogenesis. in Current Topics in Developmental Biology (ed. Ordahl, C. P.) vol. 47 81–105 (Academic Press, 1999).

32. Nowotschin, S. et al. The emergent landscape of the mouse gut endoderm at single-cell resolution. Nature 569, 361–367 (2019).

33. Lawson, K. A., & Hage, W. J. Clonal analysis of the origin of primordial germ cells in the mouse. Ciba Foundation symposium, 182, 68–91 (1994).

34. Saitou, M. & Yamaji, M. Primordial germ cells in mice. Cold Spring Harbor Perspectives in Biology 4, (2012).

35. Saitou, M., Barton, S. C. & Surani, M. A. A molecular programme for the specification of germ cell fate in mice. Nature 418, 293–300 (2002).

36. Tam, P. P. L. & Snow, M. H. L. Proliferation and migration of primordial germ cells during compensatory growth in mouse embryos. Journal of embryology and experimental morphology 64, 133–47 (1981).

37. Tanaka, Y. et al. Circulation-Independent Differentiation Pathway from Extraembryonic Mesoderm toward Hematopoietic Stem Cells via Hemogenic Angioblasts. Cell Reports 8, 31–39 (2014).

38. Rhee, J. M. et al. In vivo imaging and differential localization of lipid-modified GFP-variant fusions in embryonic stem cells and mice. genesis 44, 202–218 (2006).

39. Egli, D., Rosains, J., Birkhoff, G. & Eggan, K. Developmental reprogramming after chromosome transfer into mitotic mouse zygotes. Nature 447, 679–685 (2007).

40. Mesnard, D., Filipe, M., Belo, J. A. & Zernicka-Goetz, M. The Anterior-Posterior Axis Emerges Respecting the Morphology of the Mouse Embryo that Changes and Aligns with the Uterus before Gastrulation. Current Biology 14, 184–196 (2004).

41. Bedzhov, I., Leung, C. Y., Bialecka, M. & Zernicka-Goetz, M. In vitro culture of mouse blastocysts beyond the implantation stages. Nature Protocols 9, 2732–2739 (2014).

42. Lalit, P. A. et al. Lineage Reprogramming of Fibroblasts into Proliferative Induced Cardiac Progenitor Cells by Defined Factors. Cell stem cell 18, 354–367 (2016).

43. Zilionis, R. et al. Single-cell barcoding and sequencing using droplet microfluidics. Nature Protocols 12, 44–73 (2017).

44. Klein, A. M. et al. Droplet Barcoding for Single-Cell Transcriptomics Applied to Embryonic Stem Cells. Cell 161, 1187–1201 (2015).

45. A, B. J. et al. The dynamics of gene expression in vertebrate embryogenesis at single-cell resolution. Science 360, eaar5780 (2018).

46. Andrews, S. FastQC: a quality control tool for high throughput sequence data. (2010).

47. Parekh, S., Ziegenhain, C., Vieth, B., Enard, W. & Hellmann, I. zUMIs - A fast and flexible pipeline to process RNA sequencing data with UMIs. GigaScience 7, (2018).

48. Stuart, T. et al. Comprehensive Integration of Single-Cell Data. Cell 177, 1888–1902.e21 (2019).

49. Wolock, S. L., Lopez, R. & Klein, A. M. Scrublet: Computational Identification of Cell Doublets in Single-Cell Transcriptomic Data. Cell Systems 8, 281–291.e9 (2019).

50. Korsunsky, I. et al. Fast, sensitive and accurate integration of single-cell data with Harmony. Nature Methods 16, 1289–1296 (2019).

51. Wolf, F. A., Angerer, P. & Theis, F. J. SCANPY: large-scale single-cell gene expression data analysis. Genome Biology 19, 15 (2018).

52. Bergen, V., Lange, M., Peidli, S., Wolf, F. A. & Theis, F. J. Generalizing RNA velocity to transient cell states through dynamical modeling. Nature biotechnology 38, 1408–1414 (2020).

53. Schindelin, J. et al. Fiji: an open-source platform for biological-image analysis. Nature methods 9, 676–682 (2012).

54. Boulanger, J. et al. Patch-Based Nonlocal Functional for Denoising Fluorescence Microscopy Image Sequences. IEEE Transactions on Medical Imaging 29, 442–454 (2010).

